# GenoWAP: Post-GWAS Prioritization Through Integrated Analysis of Genomic Functional Annotation

**DOI:** 10.1101/019539

**Authors:** Qiongshi Lu, Xinwei Yao, Yiming Hu, Hongyu Zhao

## Abstract

**Motivation:** Genome-wide association study (GWAS) has been a great success in the past decade. However, significant challenges still remain in both identifying new risk loci and interpreting results. Bonferroni-corrected significance level is known to be conservative, leading to insufficient statistical power when the effect size is moderate at risk locus. Complex structure of linkage disequilibrium also makes it challenging to separate causal variants from nonfunctional ones in large haplotype blocks.

**Results:** We describe GenoWAP, a post-GWAS prioritization method that integrates genomic functional annotation and GWAS test statistics. The effectiveness of GenoWAP is demonstrated through its applications to Crohn’s disease and schizophrenia using the largest studies available, where highly ranked loci show substantially stronger signals in the whole dataset after prioritization based on a subset of samples. At the single nucleotide polymorphism (SNP) level, top ranked SNPs after prioritization have both higher replication rates and consistently stronger enrichment of eQTLs. Within each risk locus, GenoWAP is also able to distinguish functional sites from groups of correlated SNPs.

**Availability and Implementation:** GenoWAP is freely available on the web at http://genocanyon.med.yale.edu/GenoWAP

## Introduction

In the past ten years, genome-wide association studies (GWAS) have been designed and applied to identify disease genes for almost all complex diseases. As of January 15, 2015, 15,216 single nucleotide polymorphisms (SNP) from over 2,000 publications have been documented in the GWAS Catalog (Hindorff, et al., 2009). Despite its great success in identifying disease-associated loci, scientists have noted several limitations of current GWAS approaches. First, although linkage disequilibrium (LD) is the basis of GWAS, it also hinders the interpretation of association results. Due to the complex LD structure among SNPs, it is the disease-associated haplotype blocks containing hundreds of thousands of nucleotides that are identified in GWASs. Therefore, the resolution of GWAS is not sufficient for distinguishing causal variants from a large group of correlated SNPs, especially in non-coding regions where the mechanism of genomic function is still largely unknown (Cooper and Shendure, 2011; Visscher, et al., 2012; Ward and Kellis, 2012). Second, although Bonferroni-corrected significance threshold (e.g. 5×10^−8^) is widely accepted as the standard cutoff in GWAS analysis, it is well known that Bonferroni correction is too conservative when the number of hypotheses is large and there are many weak to moderate signals. In fact, for most complex diseases, numerous genomic loci are involved in disease etiology while each locus only has a moderate effect size. Therefore, studies based on high-throughput genomic scan may be underpowered if the sample size is not large enough. This has led to so-called missing heritability, which refers to the gap between the narrow-sense heritability estimated from twin/pedigree analysis and the proportion of the variance explained by significant SNPs identified from GWAS, that has been reported for many diseases (Manolio, et al., 2009; Witte, et al., 2014). One explanation of missing heritability is the insufficient statistical power to identify all the disease-associated SNPs (Eichler, et al., 2010).

Variant prioritization techniques are crucial for post-GWAS analysis on different scales. Locally, it can reveal truly functional variants within each significant locus. Globally, signals at some loci can be enhanced if proper prior information is used. Many variant prioritization methods have been proposed (Hou and Zhao, 2013). Supervised-learning-based statistical tools for predicting deleterious variants are probably the richest among available approaches. So far, most of the existing deleteriousness prediction tools only focus on protein-coding genes in the human genome. However, coding-region-based tools are not sufficient for post-GWAS prioritization because nearly 90% of the significant SNPs identified in GWAS reside in the non-coding genome (Hindorff, et al., 2009). A few tools targeting non-coding variants have been proposed (Fu, et al., 2014; Kircher, et al., 2014; Ritchie, et al., 2014; Shihab, et al., 2015). Detailed comparisons of these methods were reviewed elsewhere (Cooper and Shendure, 2011; Wang, et al., 2015). Unlike the extensively studied protein-altering variants, very few non-coding pathogenic variants have been revealed so far (Ward and Kellis, 2012). Therefore, existing non-coding variant prioritization tools based on supervised-learning may suffer from the potentially biased training data. Their performance in post-GWAS prioritization remains to be further investigated. Finally, although deleteriousness of a single SNP is crucial for identifying causal variants, it does not provide all the information needed in post-GWAS prioritization, where each SNP in GWAS also carries information of nearby variants that are not genotyped. A better informed post-GWAS prioritization method should be able to measure the functional potential for the surrounding region of each genotyped marker.

Recently, Lu et al. developed GenoCanyon, a statistical framework to predict functional non-coding regions in the human genome through integrated analysis of multiple biochemical signals and genomic conservation measures (Lu, et al., 2015). Its unsupervised-learning framework makes GenoCanyon suffer less from our limited knowledge of non-coding genome. Moreover, since the resolution of its functional prediction is at the nucleotide level, it is possible to use GenoCanyon scores to evaluate the surrounding region of each genotyped SNP. In this paper, we propose GenoWAP (Genome Wide Association Prioritizer), a post-GWAS prioritization method that integrates GenoCanyon functional prediction and GWAS p-values. We apply the method on two smaller GWASs of Crohn’s disease and schizophrenia, respectively, to prioritize SNPs. The performance is evaluated using the results from large GWAS meta-analyses of these two diseases. Compared to the top loci ranked on p-values only, top ranked loci after prioritization tend to show substantially stronger signals in large GWAS studies. Within each locus, GenoWAP is able to distinguish true signals among highly correlated SNPs. The method has the potential to reduce noises caused by LD and rescue marginal signals in GWASs with insufficient sample sizes.

## Methods

### Statistical model

For each SNP, we define *Z* to be the indicator of general functionality, and define *Z*_*D*_ to be the indicator of disease-specific functionality. More specifically, if a SNP or its surrounding region is active in any genomic functional pathway, then *Z* equals to 1. If this SNP or the surrounding region is involved in the disease pathway, then *Z*_*D*_ equals to 1. For each SNP, we use *p* to denote its p-value obtained from the standard GWAS analysis.

The goal of post-GWAS prioritization is to assign each SNP a new score that measures its importance. A reasonable quantity is the conditional probability of being disease-specific functional given the p-value, i.e. *P*(*Z*_*D*_ = 1|*p*). Using Bayes formula, we can rewrite the conditional probability as below:

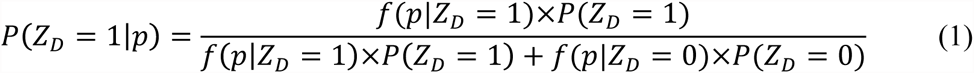

Based on the definitions of *Z* and *Z*_*D*_, we know that the SNPs satisfying *Z*_*D*_ = 1 must be a subset of the SNPs satisfying *Z* = 1. This is because if a SNP is disease-specific functional, then it has to be functional in the general sense. Therefore, we get the following formula.

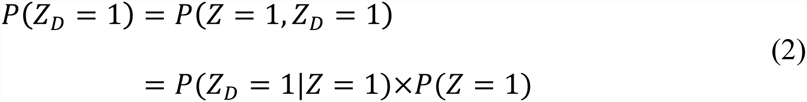

Therefore, in order to calculate the conditional probability *P*(*Z*_*D*_ = 1/*p*) for a marker, we need its prior probability of being functional, i.e. *P*(*Z* = 1); we also need the p-value density for disease-specific functional markers, i.e. *f*(*p*|*Z*_*D*_ = 1), and the p-value density for markers that are not related to the disease, i.e. *f*(*p*|*Z*_*D*_ = 0); finally, we need an estimate for the conditional probability of being disease-specific functional given the marker is functional in the general sense, i.e. *P*(*Z*_*D*_ = 1|*Z* = 1).

## Estimation

Recently, Lu et al. developed GenoCanyon, an unsupervised-learning-based statistical framework that predicts the functional potential for each nucleotide in the human genome (Lu, et al., 2015). For each SNP in our dataset, we use the mean GenoCanyon functional score of its surrounding 10,000 base pairs as the prior probability *P*(*Z* = 1). Different from using variant-based annotation tools as the prior knowledge, this prior information not only measures the importance of the genotyped marker, but also evaluates its surrounding region where the ungenotyped causal variants may reside.

Next, we partition all the SNPs into functional (Z = 1) and non-functional (Z = 0) subgroups based on the calculated mean GenoCanyon score with cutoff 0.1. Since the GenoCanyon functional score has a bimodal pattern, this partition is not sensitive to the cutoff choice. There are two major reasons why we do the partition. First, this can be viewed as a noise reduction step. After removing the non-functional markers, the signal pattern in the functional subgroup is amplified (**Figure 1**). The proportion of disease-related markers in the remaining (functional) subgroup also increased, which leads to more stable estimates in the following steps. Second, we can now empirically estimate the p-value density for non-functional markers, i.e. *f*(*p*|*Z* = 0). Since the p-values are acquired from a disease-specific case-control study, we assume that the p-values for markers that are not related to the disease should behave just like the p-values for markers that are not functional entirely. Mathematically, this assumption is characterized as the equation below.

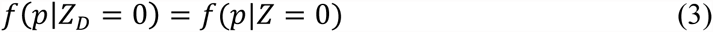

**Figure 1.**
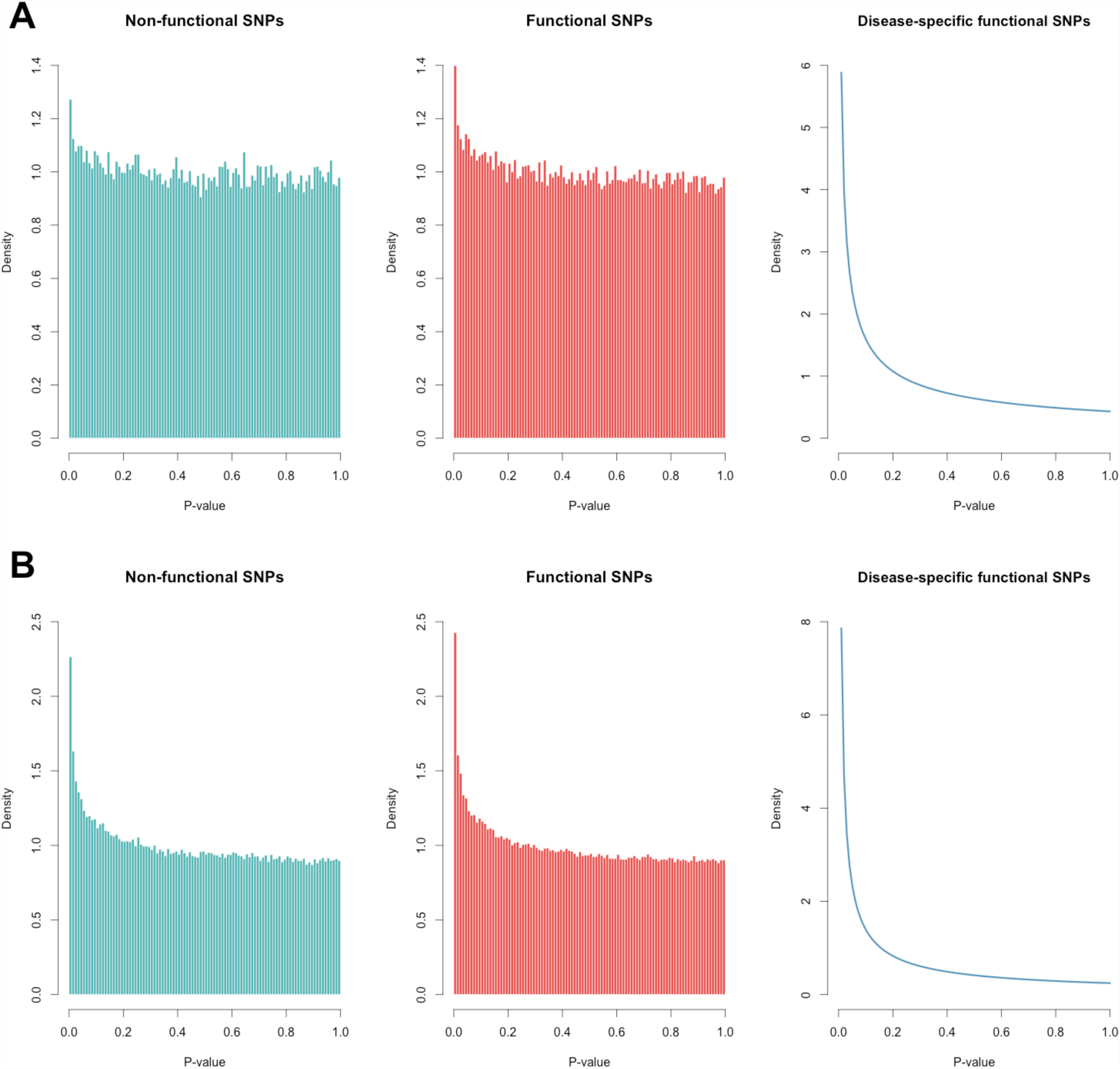
P-value densities of different subgroups of SNPs. A) P-value histogram of non-functional SNPs (Z=0, green), p-value histogram of functional SNPs (Z=1, red), and estimated p-value density of disease-specific functional SNPs (Z_D_=1, blue) in the NIDDK GWAS of Crohn’s disease. B) P-value histogram of non-functional SNPs (Z=0, green), p-value histogram of functional SNPs (Z=1, red), and estimated p-value density of disease-specific functional SNPs (Z_D_=1, blue) in the PGC2011 GWAS of schizophrenia.

Based on this assumption, we can estimate *f*(*p*|*Z*_*D*_ = 0) using the p-values for SNPs in the non-functional subgroup. Notably, it may seem natural to assume (*p*|*Z*_*D*_ = 0) follows a uniform distribution. However, the p-value of a marker with *Z*_*D*_ = 0 can actually be driven by a nearby disease-related marker due to LD. The empirically estimated density can capture a certain amount of LD information, which is complex and non-trivial to model. Moreover, it is common to see some variants with low minor allele frequencies in GWAS samples. The p-values for these markers will form a spike near 1 in the p-value density. The empirically estimated density is also able to account for this artifact. We propose to use histogram for density estimation, because it has stable performance near the boundary. In fact, the p-value boundary near 0 is where the real signals reside, and the boundary near 1 occasionally has the artifact issue caused by rare variants. Histogram is able to capture both issues. Moreover, the sample size in this framework is the number of markers, which is usually large in GWAS studies. Therefore histogram is a reasonable choice for density estimation. The number of bins is chosen based on cross-validation. It still remains to estimate the p-value density for disease-related markers *f*(*p*|*Z*_*D*_ = 1), and the conditional probability *P*(*Z*_*D*_ = 1|*Z* = 1). Now, we partition the functional subgroup (*Z* = 1) into finer subgroups. First, based on equation (3), it is straightforward to show that

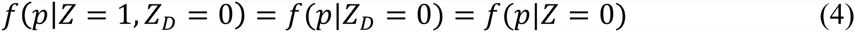

Therefore, the p-value density for functional markers is the following mixture.

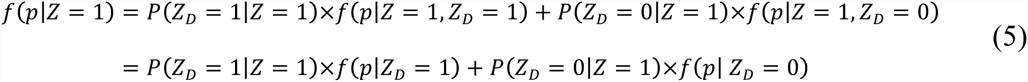

In formula (5), *f*(*p*|*Z*_*D*_ = 0) has already been estimated in previous steps. We further assume a parametric form of *f*(*p*|*Z*_*D*_ = 1). In a recent work of Chung et al., they showed that beta distribution is a robust approximation of p-value distribution under some general assumptions of SNP effect size (Chung, et al., 2014). We adopt the same assumption.

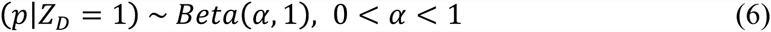

The constraint 0 < *Z* α 1 guarantees that a smaller p-value is more likely to occur than a larger p-value. Then, we apply the EM algorithm on all the p-values in the functional subgroup. One advantage of beta distribution assumption is that each iteration in the EM algorithm has a closed-form expression. In this way, we acquire the estimates for both *P*(*Z*_*D*_ = 1|*Z* = 1) and *P*(*p*|*Z*_*D*_ = 1). Then, all missing pieces in formula (1) have been estimated. We calculate the conditional probability *P*(*Z*_*D*_ = 1|*p*) for all the SNPs using these estimates. This quantity is referred to as the posterior score in this paper.

## Results

### Application to Crohn’s disease

Several GWASs of different scales have been performed for Crohn’s disease. The largest GWAS meta-analysis, which identified 71 disease-associated loci, is one of the most successful GWASs to date (Franke, et al., 2010). We applied GenoWAP on a smaller Crohn’s disease GWAS conducted by the North American National Institute of Diabetes and Digestive and Kidney Diseases (NIDDK) IBD Genetics Consortium, and tested the results using the large meta-analysis done by the International Inflammatory Bowel Disease Genetics Consortium (IIBDGC). Cohort information is listed in **Supplementary Table 1**. Details of both studies have also been reported previously (Franke, et al., 2010; Rioux, et al., 2007). It is worth noting that the samples in these two studies overlap with each other. However, the goal for this paper is not to replicate the detected signals in an independent cohort. Instead, we seek to better prioritize signals using only a small sample size. In order to test the performance, the results from the largest study available are used as the gold standard.

For each SNP in the dataset, define *p* to be the GWAS p-value, and define *Z*_*D*_ to be the indicator of disease-specific functionality. The posterior probability of being disease-specific functional, i.e. *P*(*Z*_*D*_ = 1|*p*), is used to prioritize SNPs (See **Methods**). This score will be referred to as the posterior score in the following sections. Test statistics of the NIDDK study were downloaded from dbGap (**Supplementary Table 1**). Among the 298,391 SNPs, 70 were deleted due to unavailable hg19 genomic locations. We calculated the posterior scores for the remaining 298,321 SNPs (**Supplementary Figure 1**). Test statistics of the IIBDGC meta-analysis were downloaded from the IIBDGC website (http://www.ibdgenetics.org). The dataset contains 953,241 SNPs, including 262,621 SNPs overlapping with the NIDDK dataset.

A total of 71 loci passed genome-wide significance level in the validation stage of IIBDGC meta-analysis, including 32 previously reported risk loci and 39 newly confirmed risk loci (Franke, et al., 2010). We ranked the 298,321 SNPs in the NIDDK study based on their p-values and posterior scores, respectively. Then, within each locus among the 71 loci, we compared the rank of the lowest p-value to the rank of the largest posterior score. 56 out of 71 loci (79%) had an improved rank, 3 loci (4%) had an equal rank, while only 12 loci (17%) had a reduced rank (**Supplementary Table 2**). The probability of having an increased rank is significantly higher than that of having a decreased rank (p-value = 3.11×10^−8^, one-sided binomial test).

Next, we compared the top 20 loci with the smallest p-values to the top 20 loci with the largest posterior scores in the NIDDK study. The locus information and the lowest meta-analysis p-value at each locus are listed in **Table 1**. 14 out of 20 loci are shared between the two lists. Interestingly, the posterior-specific loci, i.e. the loci that show up only in the list based on posterior score, showed substantially stronger signals in the IIBDGC meta-analysis compared to the p-value-specific loci (**Table1**, **Figure 2a**). For example, the risk locus on chromosome 10q22 was a genome-wide significant locus in the meta-analysis (rs1250550, *P*_*meta*_= 2.00×10^−10^). Although the same SNP, rs1250550, had the lowest p-value at this locus in the NIDDK dataset (*P*_*NIDDK*_ = 5.95×10^−5^), the signal was not strong enough to make this locus surpass other loci such as the one on chromosome 2q24 (rs6733000, *P*_*NIDDK*_ = *2*.01×10^−5^ **Table 1**). However, with posterior scores, locus 10q22 was ranked as the 17^th^ top locus, while the highest posterior score at locus 2q24 was only 0.0142, which agrees with its weak signal in the meta-analysis result (*P*_*meta*_ = 0.019). Overall, two posterior-specific loci were genome-wide significant in the meta-analysis, while the lowest *P*_*meta*_ among the six p-value-specific loci was only 1.10× 10^−4^. These results show that our method can effectively reduce noises likely due to LD and chance and enhance true signals at disease risk loci.

**Figure 2.**
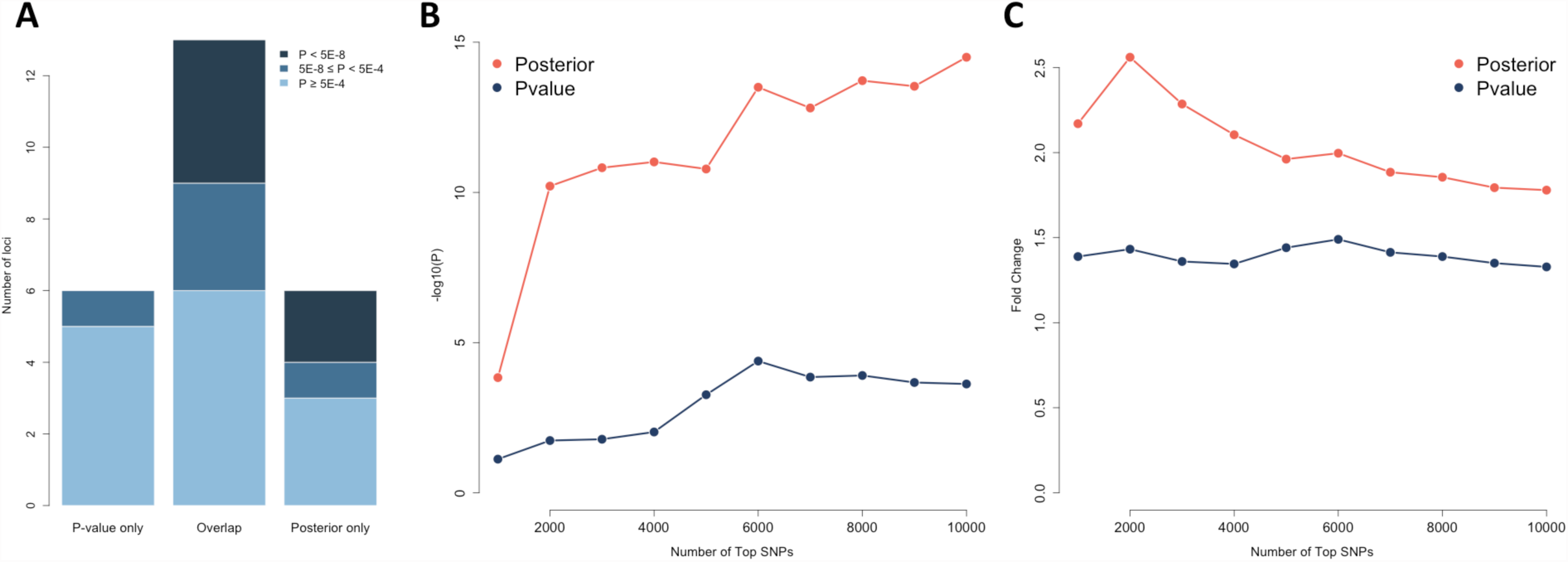
Global performance in studies of Crohn’s disease. A) Signals at p-value-specific, overlapped, and posterior-specific loci in the IIBDGC meta-analysis. The top 20 loci based on p-values in the NIDDK study are compared with the top 20 loci based on their posterior scores. Each locus is evaluated using the signal strength in the IIBDGC meta-analysis. Darker color indicates stronger signals in the meta-analysis. B) Enrichment of whole-blood eQTLs in the top SNPs selected based on p-value and posterior score. The vertical axis shows the transformed p-value of hypergeometric test. C) Fold enrichment of whole-blood eQTLs in the top SNPs selected based on p-value and posterior score. The vertical axis shows the ratio of observed and expected overlaps between eQTLs and highly ranked SNPs.

**Table 1.** The top loci with the strongest signals in the NIDDK study.

| Top 20 loci based on p-value<sup>a</sup> |  |  |  | Top 20 loci based on posterior score<sup>b</sup> |  |  |  |
| --- | --- | --- | --- | --- | --- | --- | --- |
| Chr. | Leading SNP | P<sub>NIDDK</sub> | P<sub>meta</sub> | Chr. | Leading SNP | Posterior | P<sub>meta</sub> |
| 16q12 | rs2076756 | 1.26E-14 | 4.00E-69 | 16q12 | rs2076756 | 0.999996866 | 4.00E-69 |
| 1p31 | rs7517847 | 2.99E-13 | 9.90E-65 | 1p31 | rs7517847 | 0.999972913 | 9.90E-65 |
| 2q37 | rs2241880 | 4.40E-08 | 6.70E-41 | 2q37 | rs2241880 | 0.987698022 | 6.70E-41 |
| 4p13 | rs16853571 | 5.59E-07 | 2.60E-03 | 4p13 | rs16853571 | 0.936895112 | 2.60E-03 |
| 12p13 | rs886898 | 1.05E-06 | NA<sup>c</sup> | 18q21 | rs937815 | 0.911676708 | 1.40E-05 |
| 18q21 | rs937815 | 1.88E-06 | 1.40E-05 | 12p13 | rs886898 | 0.890272531 | NA<sup>c</sup> |
| **1q23** | **rs2343331** | **2.46E-06** | **1.70E-03** | 3q23 | rs6439924 | 0.882792742 | 8.30E-04 |
| 3q23 | rs6439924 | 2.89E-06 | 8.30E-04 | 1p31 | rs2819130 | 0.773882741 | 2.20E-03 |
| **9q22** | **rs10821091** | **9.39E-06** | **5.40E-04** | 22q12 | rs4821544 | 0.752640451 | 1.80E-05 |
| 1p31 | rs2819130 | 1.23E-05 | 2.20E-03 | 11q13 | rs2712800 | 0.741211726 | 8.10E-05 |
| **14q22** | **rs1188157** | **1.23E-05** | **7.20E-04** | **9q21** | **rs4878061** | **0.697750241** | **5.20E-04** |
| 10q21 | rs224136 | 1.23E-05 | 4.40E-22 | 10q21 | rs224136 | 0.663868799 | 4.40E-22 |
| **1q23** | **rs723821** | **1.42E-05** | **1.10E-04** | **8q22** | **rs10505007** | **0.649891635** | **2.10E-04** |
| 22q12 | rs4821544 | 1.71E-05 | 1.80E-05 | **15q25** | **rs3743195** | **0.630338138** | **3.80E-03** |
| 11q13 | rs2712800 | 1.72E-05 | 8.10E-05 | 20q13 | rs4810663 | 0.626012472 | 2.40E-03 |
| 1q31 | rs2490271 | 1.88E-05 | 7.50E-03 | **7q36** | **rs4721** | **0.622452751** | **2.30E-03** |
| 8q23 | rs2044999 | 1.89E-05 | 8.40E-04 | **10q22** | **rs1250550** | **0.599609304** | **2.00E-10** |
| 20q13 | rs4810663 | 1.93E-05 | 2.40E-03 | 8q23 | rs2044999 | 0.598983918 | 8.40E-04 |
| **16q24** | **rs8050910** | **2.00E-05** | **2.40E-03** | 1q31 | rs2490271 | 0.58179581 | 7.50E-03 |
| **2p24** | **rs6733000** | **2.01E-05** | **1.90E-02** | **5p13** | **rs4613763** | **0.577762625** | **7.00E-36** |

<sup>a</sup>Top 20 loci with the smallest p-values in the NIDDK study. Loci are ordered according to the p-values. Loci in boldface are those not shared in both lists.
<sup>b</sup>Top 20 loci with the largest posterior scores in the NIDDK study. Loci are ordered according to the posterior scores. Loci in boldface are those not shared in both lists.
<sup>c</sup>The IIBDGC meta-analysis does not contain any SNP at this locus.

To see if SNPs with high posterior scores are more enriched of eQTLs, we downloaded the whole-blood eQTL data from GTEx (http://www.gtexportal.org). The top 1000 SNPs based on p-values are not statistically significantly enriched for eQTLs (p-value = 0.076; hypergeometric test; fold enrichment = 1.39), while the enrichment for the top 1000 SNPs based on posterior scores is highly significant (p-value = 1.46×10–4; fold enrichment = 2.17). The difference becomes even more drastic when using the top 2000 SNPs, with p-values 0.018 and 6.19×10^−11^ (fold enrichment 1.43 and 2.56), respectively. When the number of top SNPs increases, the posterior-based approach dominates the p-value-based approach in both enrichment p-value and fold change (**Figures 2b** and **2c**).

In order to show how our method performs locally, we chose two genome-wide significant loci from the IIBDGC meta-analysis. First, within the risk locus on chromosome 1q23, two SNPs had substantially stronger signals than others, i.e. rs2274910 (*P*_*NIDDK*_ = 4.40×10^−4^) and rs955371 (*P*_*NIDDK*_ = 4.84×10^−4^). According to the p-values, these two SNPs are undistinguishable, because the signal at rs2274910 is only slightly stronger. However, the results from the meta-analysis clearly show the existence of two SNP clusters with strong signals at this locus (**Figure 3a**). The cluster closer to gene CD244, in which rs955371 resides, actually has stronger signals than the cluster where rs2274910 is located. Interestingly, the posterior scores capture this difference between two SNPs very well. In fact, the posterior scores for rs955371 and rs2274910 are 0.272 and 0.208, suggesting rs955371 is more likely to be functional even though its p-value is larger. The second example is the risk locus on chromosome 14q35, which is one of the 12 loci with a reduced rank under the posterior scores (**Supplementary Table 2**). Signals at this locus were not strong in the NIDDK study, with the smallest p-value only at 4.70×10^−3^ (rs1959715). Moreover, the signal peak in the NIDDK study (near 88.2M) was quite far from that in the meta-analysis, which resides in genes GALC and GPR65 (**Figure 3b**). However, the posterior scores once again capture the signal pattern in the meta-analysis. Signals near 88.2M on chromosome 14 are shrunk substantially, while the SNPs in GALC and GPR65 are pushed up as the strongest signal (rs4904410). Since these SNPs have very weak signals in their p-values, the posterior score is still low (See **Methods**). This explains the reduced rank, because the p-value-based rank of rs1959715 was compared with the posterior-based rank of rs4904410. It is worth noting that the SNPs with the strongest signals in the meta-analysis, e.g. rs8005161, were either not genotyped or dropped in the quality control steps in the NIDDK study. It is reasonable to believe that the posterior scores would have had an even better performance if imputations had been done for the NIDDK dataset.

**Figure 3.**
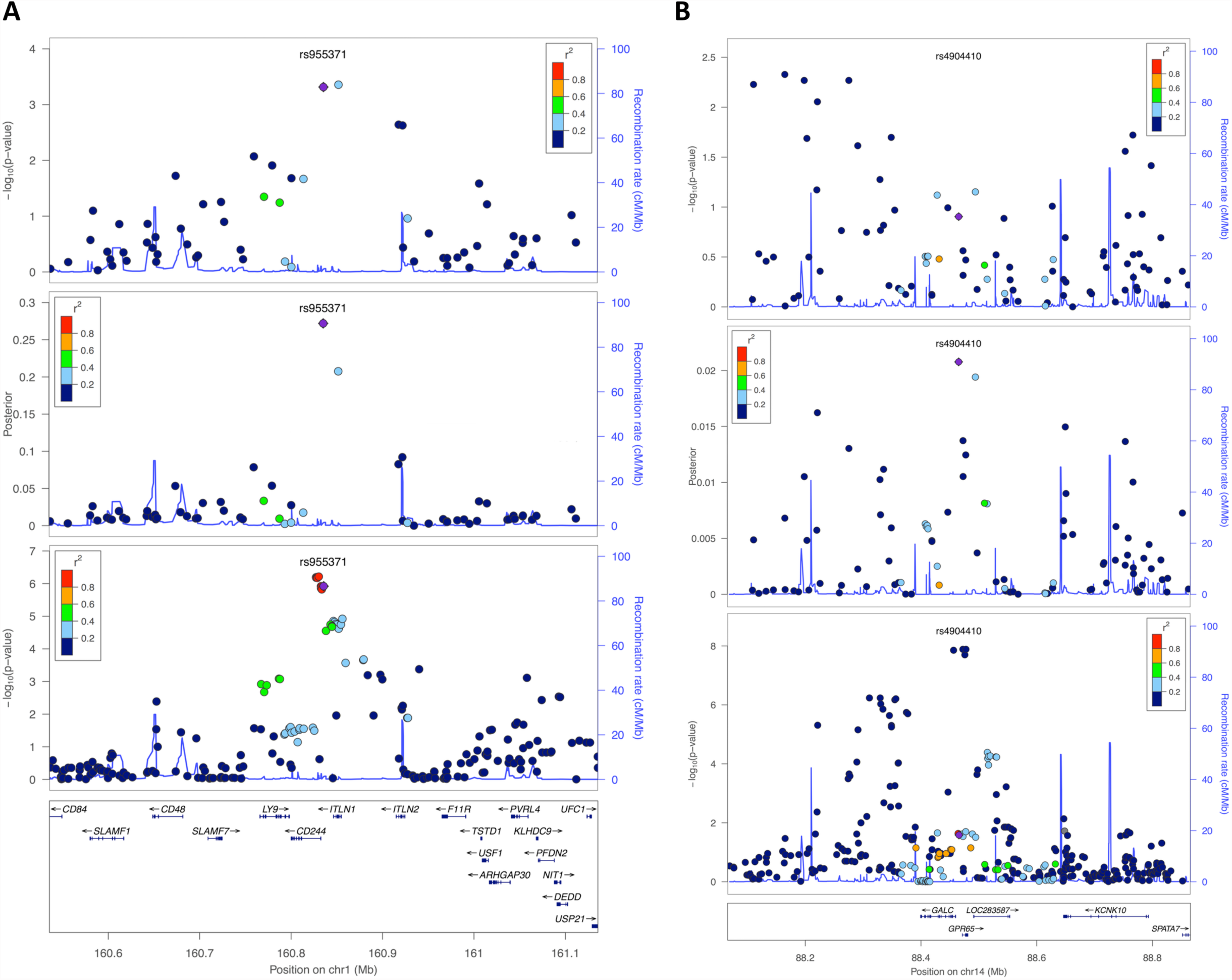
Local performance in studies of Crohn’s disease. From top to bottom, the three panels show the p-values from the NIDDK study, the posterior scores, and the p-values from the IIBDGC meta-analysis, respectively. A) Local performance at the risk locus on chromosome 1q23. The top two SNPs at this locus in the NIDDK study are undistinguishable based on their p-values. The posterior scores suggest the importance of the SNP on the left, which is in agreement with the results from the meta-analysis. B) Local performance at the risk locus on chromosome 14q35. Signals at this locus are weak in the NIDDK study, and the signal peak is different from that in the meta-analysis. The posterior score is able to pull down the noises caused by LD, and push up real signals at genes GALC and GPR65. Figures are generated using LocusZoom (Pruim, et al., 2010).

### Application to schizophrenia

In addition to Crohn’s disease, we also applied GenoWAP to schizophrenia, a major psychiatric disorder. Psychiatric Genomics Consortium (PGC), the largest international consortium in psychiatry, focuses on genetic studies of many psychiatric disorders including schizophrenia. Two large-scale GWAS mega-analyses of schizophrenia have been published. We applied GenoWAP to the earlier and smaller PGC2011 study (Consortium, 2011), and evaluated the performance using results from the larger mega-analyses published in 2014 (Consortium, 2014). Test statistics for both studies were downloaded from the PGC website (**Supplementary Table 3**). Among the 1,252,901 SNPs in PGC2011 study, 264 were removed due to unavailable hg19 locations. Posterior scores were calculated for all the remaining 1,252,637 SNPs (**Supplementary Figure 2**). PGC2014 study contains 9,444,230 SNPs, including 1,179,913 SNPs overlapping with the PGC2011 dataset.

PGC2014 study identified 108 schizophrenia-associated loci, from which we removed three loci on chromosome X because the PGC2011 dataset did not contain any SNP on sex chromosomes. We ranked the 1,252,637 SNPs in PGC2011 study based on their p-values and posterior scores, respectively. Within each locus, the rank of the lowest p-value was compared to the rank of the largest posterior score. Across the 105 loci, 68 (65%) had an improved rank, 1 locus (1%) had an equal rank, and the other 36 loci (34%) had a reduced rank (**Supplementary Table 4**). The probability of having an increased rank is significantly higher than that of having a reduced rank (p-value = 0.001, one-sided binomial test). Interestingly, among the 10 loci with the strongest signals in the PGC2014 study, 8 had an increased rank (80%). The proportion of increased or equal ranks gradually drops when more top loci in the PGC2014 study were considered, showing less confidence in weaker signals (**Supplementary Figure 3**).

Next, we compared the top 20 loci with the smallest p-values to the top 20 loci with the largest posterior scores in the PGC2011 study. In order to identify 20 independent loci, 582 SNPs were needed when using p-value as the criterion. When posterior scores were used to choose top signals, 548 SNPs were sufficient to identify 20 loci, showing better efficiency (**Figure 4a**). A total of 14 loci could be identified using both p-values and posterior scores. As for the comparisons between the 6 posterior-specific loci and the 6 p-value-specific loci, the posterior-specific loci showed better signals than the p-value-specific loci (**Table2**, **Figure 4b**) in the PGC2014 study. Four of the 6 posterior-specific loci were genome-wide significant in the PGC2014 study, whereas 2 p-value-specific loci passed the genome-wide significance level. Among the 6 p-value-specific loci, the locus on chromosome 3q26 had the strongest signal in the PGC2014 study (*P*_2014_ = 5.35× 10^−11^). This locus will be discussed in detail later.

**Figure 4.**
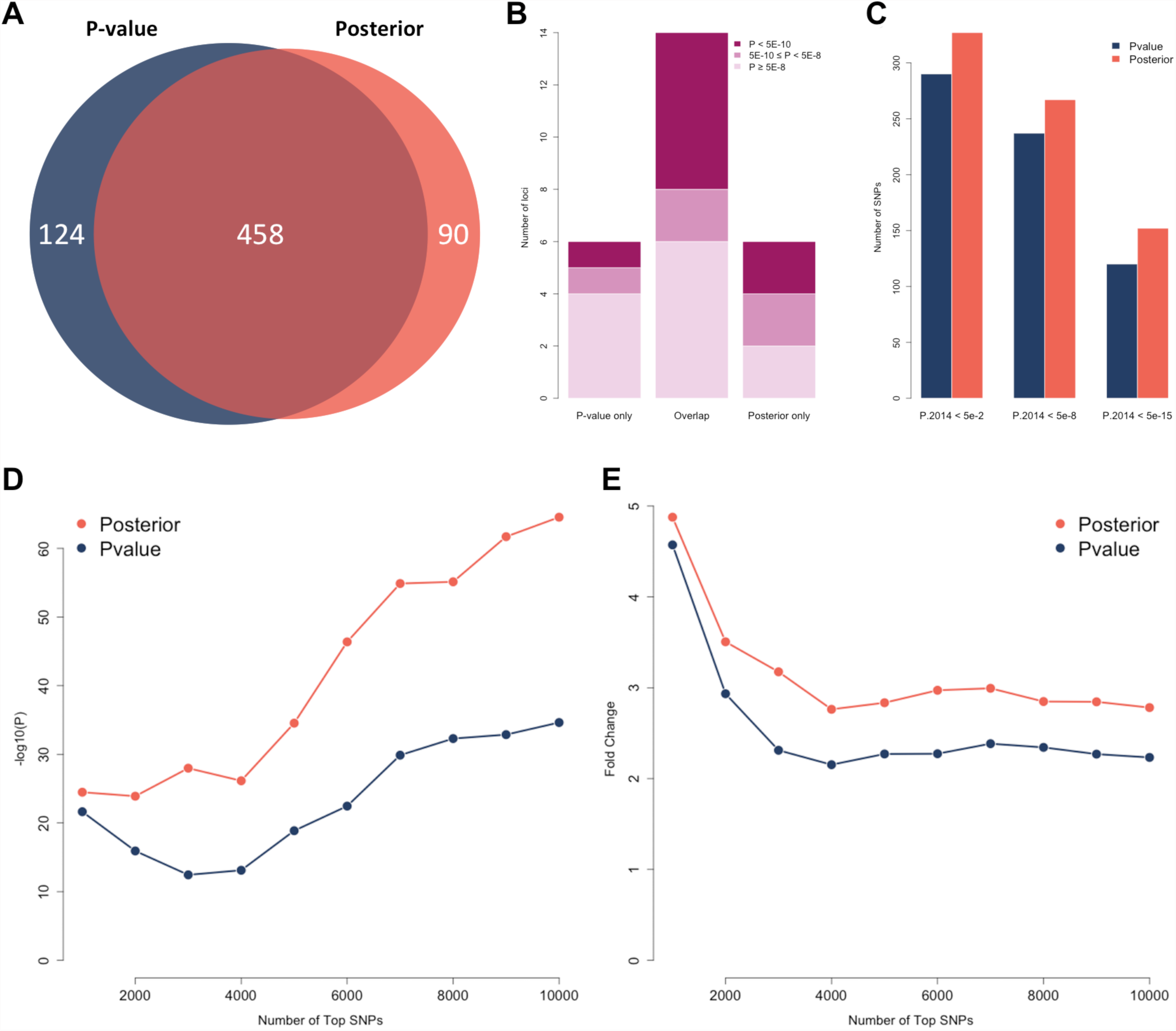
Global performance in studies of schizophrenia. A) SNPs needed for identifying 20 loci. 582 top SNPs are needed when using p-value as the criterion. 548 SNPs are sufficient when using posterior score as the criterion. B) Signals at p-value-specific, overlapped, and posterior-specific loci in the PGC2014 study. The top 20 loci based on p-values in the PGC2011 study are compared with the top 20 loci based on their posterior scores. Each locus is evaluated using the signal strength in the PGC2014 study. Darker color indicates stronger signals in the large study. C) Replication rates of SNPs before and after prioritization. The top 500 SNPs under posterior scores have substantially higher replication rates than the top 500 SNPs under p-values. D) Enrichment of whole-blood eQTLs in the top SNPs selected based on p-value and posterior score. The vertical axis shows the transformed p-value of hypergeometric test. E) Fold enrichment of whole-blood eQTLs in the top SNPs selected based on p-value and posterior score. The vertical axis shows the ratio of observed and expected overlaps between eQTLs and highly ranked SNPs.

Since imputation was done for both PGC2011 and PGC2014 studies, and the total number of SNPs is large, it is possible to compare the SNP-level replication rates when the SNPs were ranked based on p-values and posterior scores. Among the top 500 SNPs with the largest posterior scores, 327, 267, and 152 had a p-value lower than 5×10^−2^, 5×10^−8^, and 5×10^−15^ in the PGC2014 study, respectively. When choosing the top 500 SNPs based on their p-values, the corresponding numbers were 290, 237, and 120 (**Figure 4c**), respectively. A similar pattern can be observed for the top 200 SNPs (**Supplementary Table 4**). We further performed enrichment analysis for whole-blood eQTLs. The top 1000 SNPs based on the p-values were significantly enriched for eQTLs (p-value = 2.30×10^−22^, fold enrichment = 4.57), but the enrichment for the top 1000 SNPs based on the posterior scores was even stronger (p-value = 3.32×10^−25^, fold enrichment = 4.88). As the number of top SNPs increased, the posterior-based top SNPs always had stronger enrichment of eQTL than the p-value-based list (**Figures 4d** and **4e**).

Finally, we compared PGC2011 p-values, PGC2011 posterior scores, and PGC2014 p-values at two loci to further illustrate the performance of our method. The first locus is on chromosome 3q26. It had the strongest signal in PGC2014 among the p-value-specific top 20 loci (**Table 2**, *P*_2014_ = 5.35×10-11). Based on the p-values in the PGC2011 study, the strongest signals reside in the intergenic region upstream of FXR1. But the posterior scores brought down those intergenic SNPs, and enhanced the signals in FXR1 instead, which is in agreement with the results from PGC2014 (**Figure 5a**). In fact, from the PGC2014 p-values, we can clearly see that the strongest signals reside in FXR1 while the significant results for the SNPs upstream or downstream of FXR1 are likely due to LD. The second example is on chromosome 8q21 (**Figure 5b**). In the PGC2011 study, the strongest signal at this locus resides in the intergenic region between 89.7M and 89.8M. However, posterior scores removed most of the correlated SNPs at this locus, leaving three separate peaks as candidate functional spots. The first peak lies right upstream of MMP16. The second peak is more upstream (∼89.6M), and is suggested to be the strongest signal source. The SNPs with the lowest p-values in PGC2011 remained as a signal peak, but their posterior scores were not as strong as the peak in the middle. Most interestingly, the results from the posterior scores perfectly matched the signal patterns in the PGC2014 study. From the lowest panel in **Figure 5b**, we can clearly see two separate peaks at the same locations suggested by the posterior scores, with the one near 89.6M being the strongest signal source. Also, the SNPs between 89.7M and 89.8M had weaker signals than the peak in the middle. Notably, this entire risk locus resides in an intergenic region. This example shows that our method can effectively prioritize SNPs in the non-coding genome.

**Figure 5.**
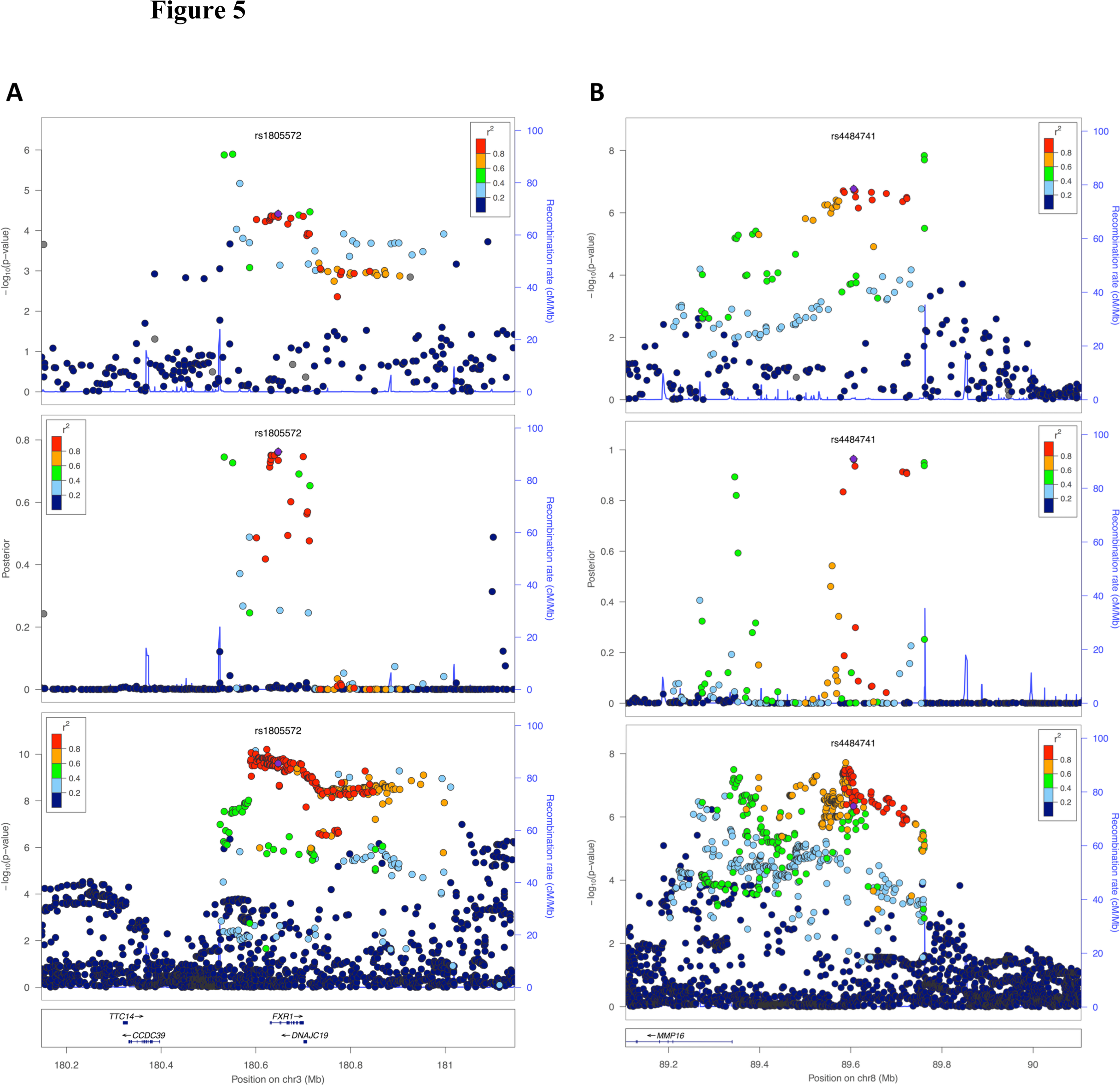
Local performance in studies of schizophrenia. From top to bottom, the three panels show the p-values from the PGC2011 study, the posterior scores, and the p-values from the PGC2014 study, respectively. A) Local performance at the risk locus on chromosome 3q26. The top signals at this locus in the PGC2011 study reside upstream of gene FXR1, while the posterior scores pull down those signals and suggest the importance of SNPs in FXR1. This agrees with the signal pattern in the PGC2014 study. B) Local performance at the risk locus on chromosome 8q21. Posterior scores diminish most of the correlated SNPs at this locus, leaving three separate signal peaks. The peak near 89.6M is suggested to be the strongest signal source, which cannot be seen using p-values from the PGC2011 study. The signal peaks suggested by posterior scores perfectly match the strongest signals in the PGC2014 study. Figures are generated using LocusZoom (Pruim, et al., 2010).

**Table 2.** The top loci with the strongest signals in the PGC2011 study.

| Top 20 loci based on p-value<sup>a</sup> |  |  |  | Top 20 loci based on posterior score<sup>b</sup> |  |  |  |
| --- | --- | --- | --- | --- | --- | --- | --- |
| Chr. | Leading SNP | P<sub>2011</sub> | P<sub>2014</sub> | Chr. | Leading SNP | Posterior | P<sub>2014</sub> |
| 6p22 | rs2021722 | 4.30E-11 | 3.86E-32 | 6p22 | rs2021722 | 0.999990069 | 3.86E-32 |
| 8q21 | rs7004633 | 1.45E-08 | 1.90E-08 | 10q24 | rs11191580 | 0.998895413 | 9.24E-18 |
| 10q24 | rs11191580 | 2.23E-08 | 9.24E-18 | 18q21 | rs17512836 | 0.998854457 | 9.09E-13 |
| 18q21 | rs17512836 | 2.35E-08 | 9.09E-13 | 11q24 | rs548181 | 0.998628374 | 2.87E-05 |
| 11q24 | rs548181 | 2.91E-08 | 2.87E-05 | 7p22 | rs1107592 | 0.99673362 | 6.12E-14 |
| 7p22 | rs10226475 | 5.06E-08 | 6.12E-14 | 15q15 | rs1869901 | 0.991093414 | 4.92E-08 |
| **8p23** | **rs10503256** | **1.96E-07** | **2.69E-08** | 1p21 | rs1625579 | 0.98743964 | 2.79E-17 |
| 3p14 | rs11130874 | 2.09E-07 | 7.68E-05 | 3p14 | rs191558 | 0.987169485 | 7.68E-05 |
| 15q15 | rs1869901 | 3.49E-07 | 4.92E-08 | 14q13 | rs10135277 | 0.986756984 | 1.52E-07 |
| 14q13 | rs10135277 | 5.11E-07 | 1.52E-07 | 9p24 | rs12352353 | 0.986087806 | 3.32E-04 |
| 1p21 | rs1625579 | 5.72E-07 | 2.79E-17 | 12p13 | rs4765905 | 0.980692241 | 2.63E-17 |
| 9p24 | rs12352353 | 6.57E-07 | 3.32E-04 | 6p21 | rs9462875 | 0.966743496 | 9.61E-07 |
| **2q31** | **rs17180327** | **6.80E-07** | **5.95E-06** | **3p21** | **rs2239547** | **0.96441905** | **3.96E-11** |
| 12p13 | rs7972947 | 7.77E-07 | 2.63E-17 | 8q21 | rs4484741 | 0.96417159 | 1.90E-08 |
| **10q26** | **rs1025641** | **8.28E-07** | **2.21E-04** | **2q37** | **rs2675968** | **0.958718346** | **3.15E-12** |
| 2q37 | rs13025591 | 1.07E-06 | 3.84E-05 | 2q37 | rs13025591 | 0.955105149 | 3.84E-05 |
| **11q22** | **rs2509843** | **1.10E-06** | **1.24E-04** | **1p36** | **rs2252865** | **0.939171332** | **2.03E-09** |
| **3q26** | **rs1879248** | **1.27E-06** | **5.35E-11** | **1q43** | **rs10803133** | **0.937014805** | **4.40E-09** |
| 6p21 | rs9462875 | 1.46E-06 | 9.61E-07 | **21q22** | **rs11702343** | **0.934578026** | **8.04E-05** |
| **11p15** | **rs4356203** | **1.90E-06** | **8.01E-06** | **22q12** | **rs9621795** | **0.925914726** | **1.26E-03** |

<sup>a</sup>Top 20 loci with the smallest p-values in the PGC2011 study. Loci are ordered according to the p-values. Loci in boldface are those not shared in both lists.
<sup>b</sup>Top 20 loci with the largest posterior scores in the PGC2011 study. Loci are ordered according to the posterior scores. Loci in boldface are those not shared in both lists.

## Discussion

In this study, we developed and applied GenoWAP to two sets of GWAS data to illustrate its performance in post-GWAS prioritization. Compared to p-values, GenoWAP posterior scores can better prioritize SNPs in many different ways. At the locus level, posterior score is more efficient in the sense that fewer SNPs are needed to identify the same number of top loci. Moreover, noises due to chance are effectively reduced, and the highly ranked loci using posterior score are more likely to be functional than the top loci selected purely based on p-values. At the SNP level, markers with high posterior scores have both better replication rates and consistently stronger enrichment of eQTLs than the top SNPs based on p-values. More importantly, within each risk locus identified in GWAS, posterior scores can effectively suggest real signals among a large number of correlated SNPs.

The performance of GenoWAP depends on the accuracy of functional annotation and the quality of GWAS data. Due to our limited understanding of non-coding genome, it is challenging to provide accurate genomic functional annotation. GenoCanyon is the first functional prediction tool at the nucleotide level. When more accurate or tissue-specific functional annotation becomes available in the future, the performance of GenoWAP may be further improved. On the other hand, GenoWAP does not play magic. If no information is contained in the GWAS dataset, then GenoWAP can only provide very limited insight.

More than 2,000 GWASs have been published in the past decade, and the number continues to grow. It is well known that our ability to identify new risk loci for complex diseases has surpassed our ability to interpret the results. However, although we are overwhelmed by the large amount of information detected in GWASs, evidence such as missing heritability still suggests that many risk loci remain to be discovered. Therefore, there is pressing need for post-GWAS prioritization tools and our method has great potential for future application. Since GenoWAP uses only p-values as the input, it is convenient to apply our method on published results, which may help reveal truly functional variants within large haplotype blocks, and ultimately help understand disease etiology. Moreover, for multi-stage GWASs, GenoWAP can be used to better prioritize SNPs from the discovery stage to the validation planning and increase the replication rates. Finally, next-generation sequencing is widely recognized as the future of genomic epidemiology. However, the high cost of sequencing usually leads to insufficient sample sizes and many other challenging issues (Sboner, et al., 2011). The combination of GenoWAP and the rich collection of publicly available GWAS data have the potential to provide functional candidates and guide sequencing analysis in the future.

## Authors’ contributions

QL conceived and wrote the original manuscript. XY, QL, and YH developed the GenoWAP software. HZ advised on genetic and statistical issues.

## Acknowledgements

We would like to thank Dr. Katerina Kechris and all the members in the Data Integration – COPD working group at SAMSI for their advice and useful discussions on this work. This study was supported by the National Institutes of Health grants R01 GM59507 and U01 HG005718, the VA Cooperative Studies Program of the Department of Veterans Affairs, Office of Research and Development, and the Yale World Scholars Program sponsored by the China Scholarship Council.

## Supplementary Information

**Supplementary Figure 1.**
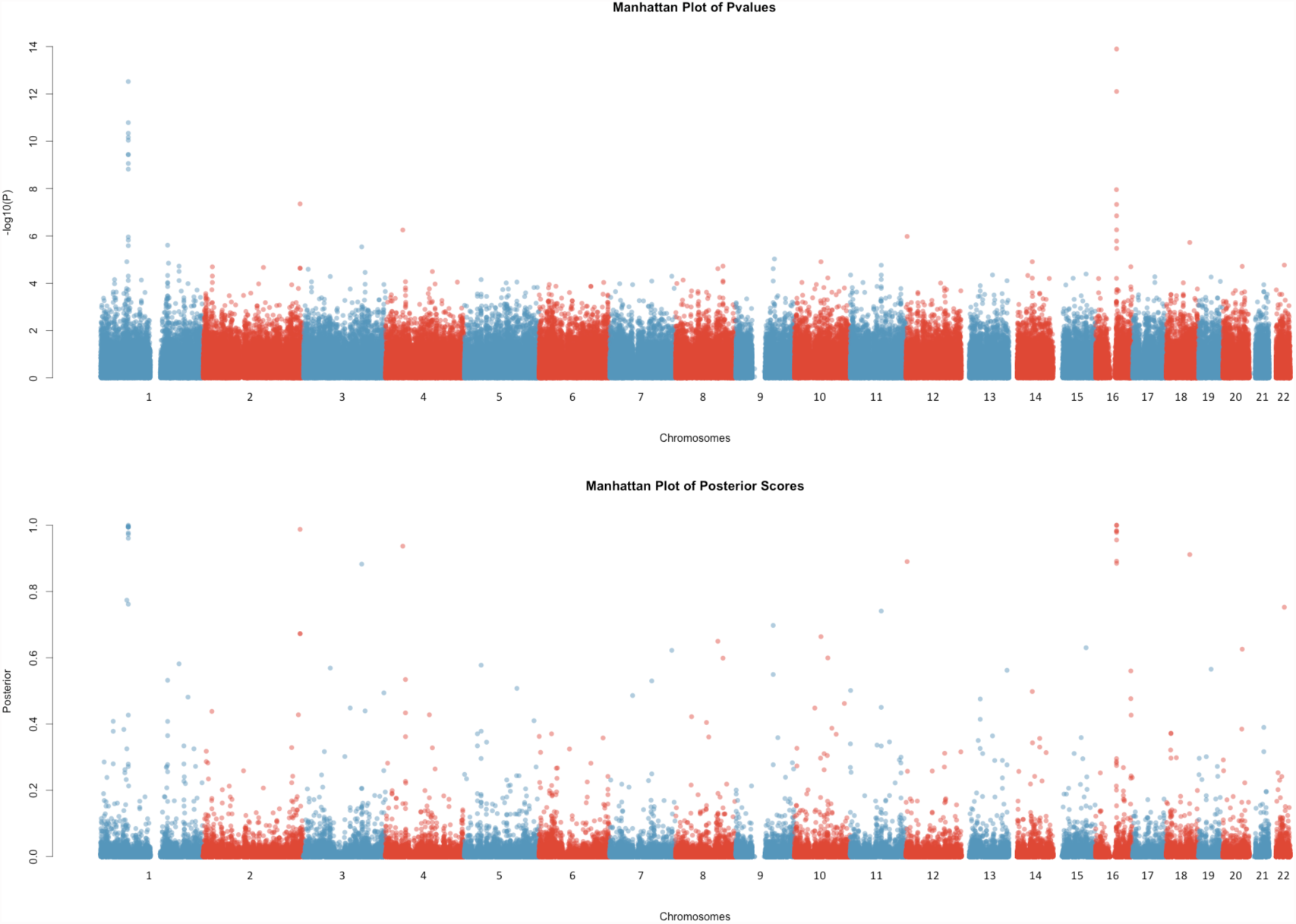
Manhattan plot of p-values and posterior scores for the NIDDK study.

**Supplementary Figure 2.**
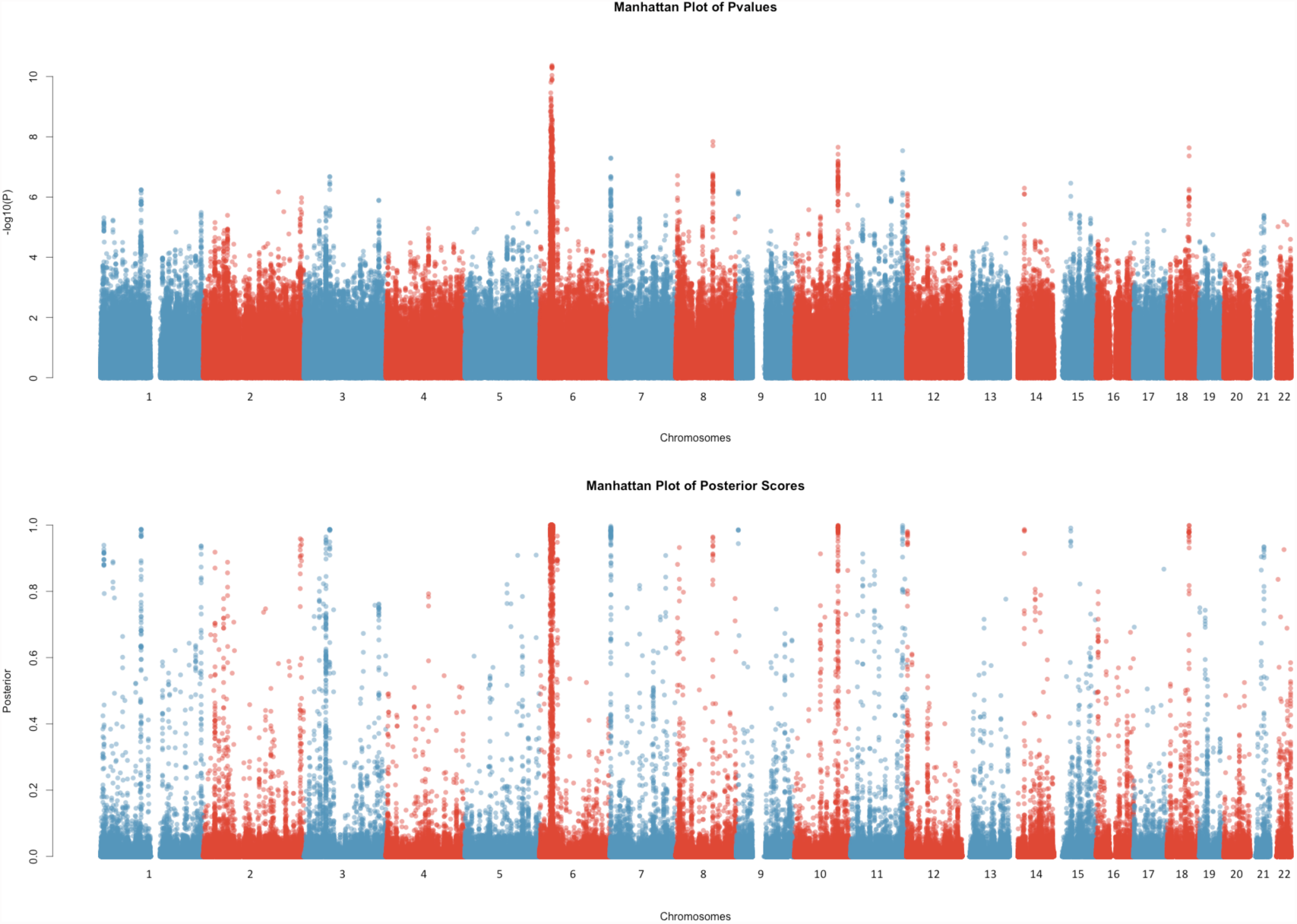
Manhattan plot of p-values and posterior scores for the PGC2011 study.

**Supplementary Figure 3.**
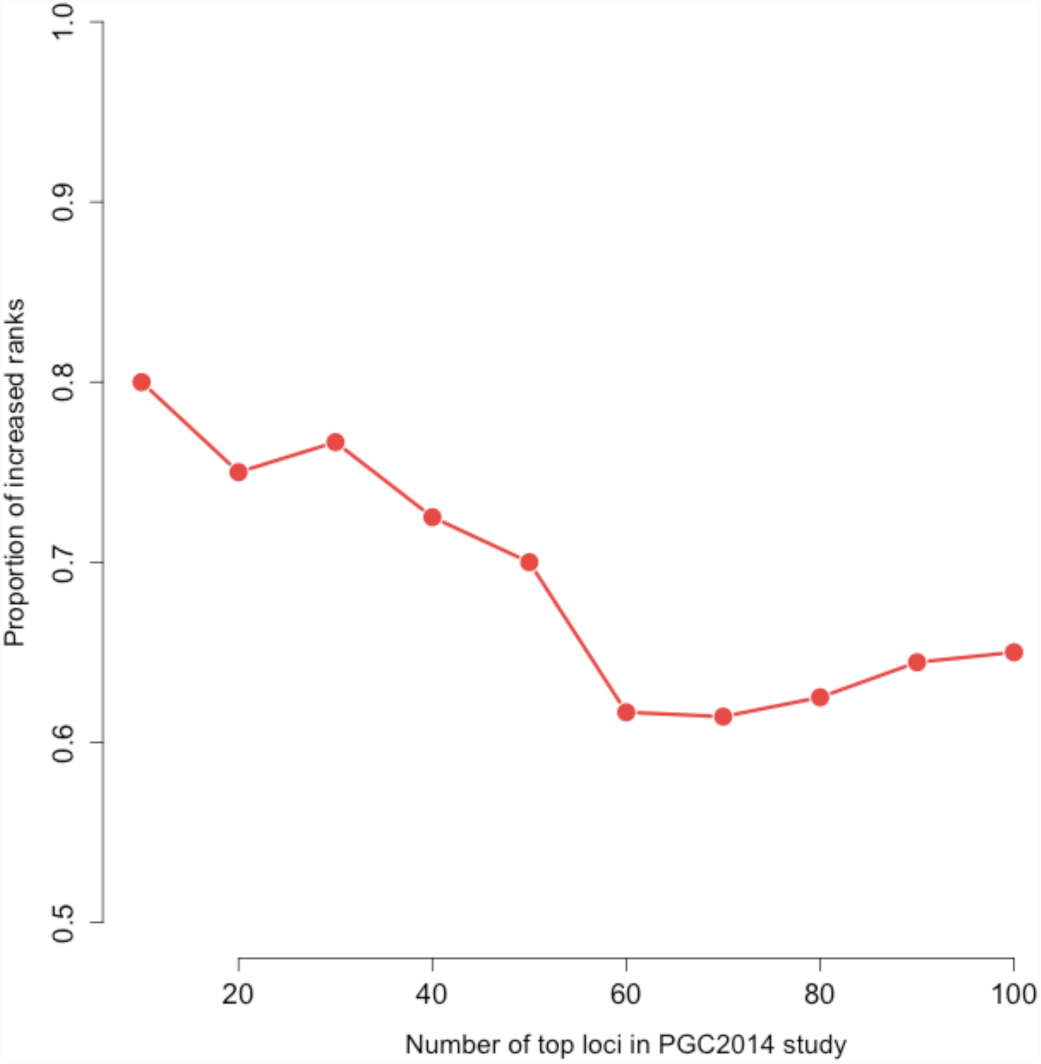
Negative association between the proportion of increased ranks under posterior score and the number of top loci being considered.

**Supplementary Figure 4.**
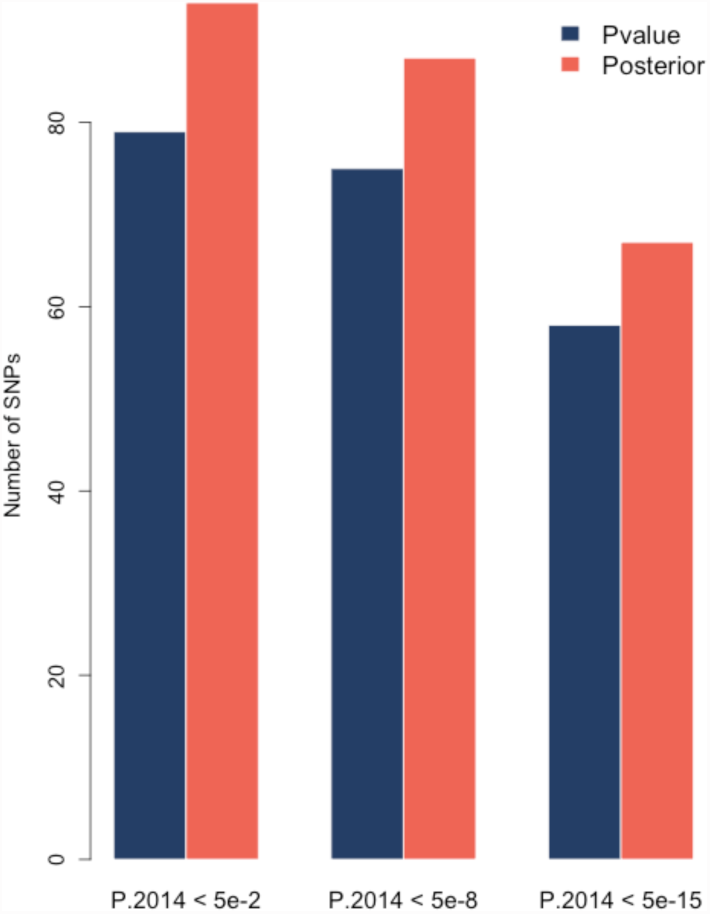
Enhanced replication rates after prioritization among the top 200 SNPs.

**Supplementary Table 1.** Basic information for NIDDK study and IIBDGC Meta-analysis.

|  | **NIDDK Study<sup>a</sup>** | **IIBDGC Meta-Analysis<sup>b</sup>** |
| --- | --- | --- |
| **Disease** | Crohn's disease | Crohn's disease |
| **# Cases** | 968<sup>c</sup> | 6,333 |
| **# Controls** | 995 | 15,056 |
| **Genotyping Platform** | Illumina HumanHap300 | Multiple |
| **# SNPs** | 298,391 | 953,241 |
| **Imputation** | No imputation | HAPMAP3 |

<sup>a</sup> The SNP-level summary statistics of the NIDDK study was downloaded from dbGap (http://www.ncbi.nlm.nih.gov/projects/gap/cgi-bin/study.cgi?study_id=phs000130.v1.p1).
<sup>b</sup> The SNP-level summary statistics of the IIBDGC meta-analysis was downloaded from the IIBDGC website (http://www.ibdgenetics.org).
<sup>c</sup> The sample size information might be inconsistent with the original publication. The case/control sample size here was extracted from the header lines of the analyses file downloaded from dbGaP.

**Supplementary Table 2.** Ranks of top signals under p-value and posterior score at 71 genome-wide significant loci of Crohn’s disease.

| Rankings based on p-values |  |  |  | Ranking based on posterior scores |  |  |
| --- | --- | --- | --- | --- | --- | --- |
| Chr. | Leading SNP | Pvalue | Rank<sup>a</sup> | Leading SNP | Posterior | Rank<sup>b</sup> |
| *32 previously confirmed Crohn's disease risk loci* |  |  |  |  |  |  |
| 1p31 | rs7517847 | 2.99E-13 | 2 | rs7517847 | 0.999972913 | 3 |
| 1p13 | rs971173 | 4.44E-03 | 1954 | rs971173 | 0.114306737 | 679 |
| 1q23 | rs2274910 | 4.40E-04 | 273 | rs955371 | 0.272006962 | 154 |
| 1q24 | rs9286879 | 8.39E-03 | 3444 | rs9286879 | 0.072577451 | 1507 |
| 1q32 | rs2297909 | 3.52E-03 | 1582 | rs12122721 | 0.104187914 | 796 |
| 2q37 | rs2241880 | 4.40E-08 | 13 | rs2241880 | 0.987698022 | 9 |
| 3p21 | rs7629936 | 4.69E-04 | 282 | rs7629936 | 0.316858359 | 111 |
| 5p13 | rs4613763 | 6.97E-05 | 68 | rs4613763 | 0.577762625 | 38 |
| 5q31 | rs2243300 | 8.76E-04 | 491 | rs2243300 | 0.243962672 | 184 |
| 5q33 | rs2112637 | 1.23E-03 | 635 | rs2112637 | 0.183061328 | 283 |
| 5q33 | rs10045431 | 1.03E-03 | 561 | rs6556377 | 0.056512976 | 2330 |
| 6p22 | rs6921781 | 8.76E-03 | 3578 | rs6921781 | 0.071176737 | 1554 |
| 6p21 | rs630379 | 1.62E-03 | 781 | rs630379 | 0.186331957 | 274 |
| 6q21 | rs2859307 | 5.53E-03 | 2373 | rs2859307 | 0.101481811 | 842 |
| 6q27 | rs9347189 | 2.69E-03 | 1230 | rs9347189 | 0.104526433 | 788 |
| 7p12 | rs2045369 | 1.44E-02 | 5604 | rs10251980 | 0.035266352 | 5016 |
| 8q24 | rs2124036 | 2.28E-02 | 8398 | rs2124036 | 0.048432287 | 2942 |
| 9p24 | rs2150192 | 3.47E-03 | 1557 | rs2150192 | 0.128407135 | 557 |
| 9q32 | rs10817694 | 1.39E-02 | 5418 | rs10817694 | 0.063059551 | 1957 |
| 10p11 | rs2504246 | 1.01E-03 | 550 | rs2492448 | 0.040627018 | 3967 |
| 10q21 | rs224136 | 1.23E-05 | 30 | rs224136 | 0.663868799 | 30 |
| 10q24 | rs888208 | 3.05E-04 | 201 | rs888208 | 0.369345672 | 82 |
| 11q13 | rs1892954 | 9.91E-04 | 543 | rs6592651 | 0.130472283 | 537 |
| 12q12 | rs545385 | 6.17E-04 | 358 | rs1444204 | 0.044793579 | 3365 |
| 13q14 | rs583271 | 1.32E-02 | 5153 | rs3764147 | 0.048506448 | 2934 |
| 16q12 | rs2076756 | 1.26E-14 | 1 | rs2076756 | 0.999996866 | 1 |
| 17q21 | rs931992 | 6.32E-03 | 2687 | rs931992 | 0.094628434 | 945 |
| 17q21 | rs9252 | 4.61E-02 | 16050 | rs9252 | 0.032894817 | 5790 |
| 18p11 | rs9303778 | 2.88E-03 | 1305 | rs9303778 | 0.14135767 | 460 |
| 19p13 | rs12977033 | 5.45E-04 | 324 | rs12977033 | 0.296653851 | 127 |
| 21q21 | rs1736148 | 2.69E-03 | 1230 | rs1736148 | 0.145846532 | 435 |
| 21q22 | rs3827246 | 1.14E-02 | 4497 | rs3827246 | 0.07032351 | 1585 |
| *39 Crohn's disease risk loci newly confirmed in the meta-analysis* |  |  |  |  |  |  |
| 1p36 | rs707455 | 4.26E-04 | 264 | rs707455 | 0.190027644 | 264 |
| 1q22 | rs6427128 | 1.72E-03 | 833 | rs6427128 | 0.181139121 | 288 |
| 1q31 | rs10922341 | 2.41E-02 | 8834 | rs12568860 | 0.012105168 | 36458 |
| 1q32 | rs4845119 | 1.07E-02 | 4287 | rs6677934 | 0.046490815 | 3156 |
| 2p23 | rs658414 | 1.04E-03 | 564 | rs517403 | 0.12858641 | 555 |
| 2p23 | rs780094 | 3.54E-02 | 12585 | rs2304681 | 0.026501082 | 8879 |
| 2p21 | rs3901678 | 2.49E-03 | 1150 | rs3901678 | 0.088740493 | 1060 |
| 2p16 | rs13003464 | 2.32E-03 | 1085 | rs13003464 | 0.128992257 | 549 |
| 2q12 | rs917997 | 1.55E-03 | 751 | rs13015714 | 0.124198461 | 589 |
| 2q33 | rs700646 | 3.90E-03 | 1748 | rs770657 | 0.02803119 | 8052 |
| 2q37 | rs16827412 | 2.69E-02 | 9748 | rs7426302 | 0.038987848 | 4243 |
| 3p24 | rs6792314 | 8.60E-05 | 79 | rs9881034 | 0.105980098 | 773 |
| 5q13 | rs7702331 | 1.28E-02 | 5005 | rs638333 | 0.049480331 | 2826 |
| 5q15 | rs4869151 | 9.50E-03 | 3857 | rs4869151 | 0.077020627 | 1358 |
| 5q31 | rs445310 | 6.52E-03 | 2764 | rs445310 | 0.094057648 | 957 |
| 5q35 | rs359457 | 6.94E-04 | 399 | rs359457 | 0.270517728 | 156 |
| 6p25 | rs11242859 | 4.68E-04 | 281 | rs11242859 | 0.314623689 | 114 |
| 6q15 | rs6939786 | 5.60E-03 | 2397 | rs6939786 | 0.099185683 | 882 |
| 6q25 | rs7746447 | 2.39E-03 | 1110 | rs7746447 | 0.154569531 | 395 |
| 8q24 | rs3922389 | 3.27E-02 | 11697 | rs3922389 | 0.03981035 | 4091 |
| 9q34 | rs4077515 | 7.33E-04 | 414 | rs4077515 | 0.264543061 | 164 |
| 10p15 | rs4750000 | 4.27E-04 | 265 | rs4750000 | 0.326925509 | 104 |
| 10q21 | rs11005962 | 4.46E-03 | 1960 | rs1416764 | 0.070168232 | 1591 |
| 10q22 | rs1250550 | 5.95E-05 | 60 | rs1250550 | 0.599609304 | 35 |
| 11q12 | rs7947046 | 1.75E-03 | 851 | rs7947046 | 0.16782237 | 338 |
| 11q13 | rs645078 | 3.61E-03 | 1615 | rs645078 | 0.116891493 | 653 |
| 13q14 | rs1900448 | 1.56E-02 | 6030 | rs1900448 | 0.059300765 | 2172 |
| 14q24 | rs174213 | 3.52E-03 | 1582 | rs174213 | 0.076308497 | 1376 |
| 14q35 | rs1959715 | 4.70E-03 | 2057 | rs4904410 | 0.020775196 | 13706 |
| 15q22 | rs745103 | 3.36E-04 | 219 | rs745103 | 0.359154463 | 88 |
| 16p11 | rs151229 | 2.18E-02 | 8101 | rs151229 | 0.025564195 | 9471 |
| 17q12 | rs991804 | 3.37E-03 | 1519 | rs991804 | 0.066547951 | 1762 |
| 19p13 | rs6511696 | 4.76E-03 | 2077 | rs3181049 | 0.049651701 | 2809 |
| 19q13 | rs2287882 | 4.49E-03 | 1973 | rs17760633 | 0.089213415 | 1051 |
| 19q13 | rs485186 | 1.95E-03 | 931 | rs504963 | 0.102902888 | 816 |
| 20q13 | rs3810481 | 1.36E-02 | 5310 | rs3795149 | 0.062480703 | 1995 |
| 22q11 | rs2298428 | 8.13E-04 | 462 | rs2298428 | 0.253181625 | 175 |
| 22q12 | rs9621049 | 2.94E-04 | 196 | rs9621049 | 0.2417048 | 190 |
| 22q13 | rs9607601 | 5.20E-04 | 307 | rs54211 | 0.151045064 | 412 |

<sup>a</sup> rank of the SNP with the smallest p-value
<sup>b</sup> rank of the SNP with the largest posterior score

**Supplementary Table 3.** Basic information for PGC2011 study and PGC2014 study.

|  | **PGC 2011 Study<sup>a</sup>** | **PGC 2014 Study<sup>a</sup>** |
| --- | --- | --- |
| **Study Type** | GWAS Mega-Analysis | GWAS Mega-Analysis |
| **Disease** | Schizophrenia | Schizophrenia |
| **# Cases** | 9,394 | 34,241<sup>b</sup> |
| **# Controls** | 12,462 | 45,604 |
| **Genotyping Platform** | Multiple | Multiple |
| **# SNPs** | 1,252,901 | 9,444,230 |
| **Imputation** | HAPMAP3 | 1000 Genomes Project |

<sup>a</sup> The SNP-level summary statistics of both studies were downloaded from the PGC website (http://www.med.unc.edu/pgc/).
<sup>b</sup> These are the number of case-control samples. Besides this, 1,235 parent affected-offspring trios were also included in this study.

**Supplementary Table 4.** Ranks of top signals under p-value and posterior score at 105 genome-wide significant loci of schizophrenia.

| Chr. | Start | Stop | Rankings based on p-values |  |  | Ranking based on posterior scores |  |  | PGC2014 Rank<sup>c</sup> |
| --- | --- | --- | --- | --- | --- | --- | --- | --- | --- |
|  |  |  | Leading SNP | Pvalue | Rank<sup>a</sup> | Leading SNP | Posterior | Rank<sup>b</sup> |  |
| 1 | 2,372,401 | 2,402,501 | rs10910078 | 2.84E-03 | 12084 | rs10910078 | 0.105625533 | 6249 | 48 |
| 1 | 8,411,184 | 8,638,984 | rs2252865 | 4.90E-06 | 714 | rs2252865 | 0.939171332 | 516 | 51 |
| 1 | 30,412,551 | 30,437,271 | rs1009080 | 5.84E-06 | 746 | rs1009080 | 0.889510826 | 636 | 59 |
| 1 | 44,029,384 | 44,128,084 | rs11210896 | 4.84E-03 | 17544 | rs3001723 | 0.054816721 | 12702 | 44 |
| 1 | 73,766,426 | 73,991,366 | rs11210205 | 1.25E-03 | 7070 | rs11210274 | 0.004268122 | 412743 | 20 |
| 1 | 97,792,625 | 98,559,084 | rs1625579 | 5.72E-07 | 395 | rs1625579 | 0.98743964 | 242 | 2 |
| 1 | 149,998,890 | 150,242,490 | rs16835254 | 1.05E-04 | 1981 | rs7521783 | 0.587006577 | 1211 | 46 |
| 1 | 177,247,821 | 177,300,821 | rs1883243 | 2.20E-02 | 53656 | rs16851048 | 0.018150739 | 59742 | 98 |
| 1 | 207,912,183 | 208,024,083 | rs2796267 | 3.81E-04 | 3724 | rs2796267 | 0.370349774 | 2010 | 99 |
| 1 | 243,503,719 | 244,002,945 | rs6703335 | 3.26E-06 | 654 | rs6703335 | 0.930827436 | 534 | 62 |
| 2 | 57,943,593 | 58,502,192 | rs11682175 | 4.07E-06 | 679 | rs2683634 | 0.88791048 | 640 | 27 |
| 2 | 72,357,335 | 72,368,185 | rs2241057 | 2.76E-02 | 64075 | rs2241057 | 0.032173977 | 27702 | 72 |
| 2 | 146,416,922 | 146,441,832 | rs2381759 | 3.66E-03 | 14379 | rs2381759 | 0.004539113 | 377737 | 56 |
| 2 | 149,390,778 | 149,520,178 | rs12614977 | 9.92E-03 | 29507 | rs12614977 | 0.047367587 | 15941 | 84 |
| 2 | 162,798,555 | 162,910,255 | rs4664442 | 7.99E-05 | 1738 | rs2052400 | 0.507027988 | 1434 | 102 |
| 2 | 185,601,420 | 185,785,420 | rs1344706 | 1.84E-04 | 2629 | rs2369595 | 0.027291935 | 34285 | 17 |
| 2 | 193,848,340 | 194,028,340 | rs17662626 | 3.09E-06 | 645 | rs17662626 | 0.007332516 | 203243 | 75 |
| 2 | 198,148,577 | 198,835,577 | rs8539 | 9.97E-04 | 6166 | rs8539 | 0.22157605 | 3253 | 30 |
| 2 | 200,161,422 | 200,309,252 | rs4673339 | 9.31E-02 | 166270 | rs4673339 | 0.009291762 | 146143 | 74 |
| 2 | 200,715,237 | 200,848,037 | rs11694369 | 2.15E-03 | 9958 | rs1509835 | 0.086500653 | 7519 | 10 |
| 2 | 225,334,096 | 225,467,796 | rs2047134 | 1.35E-02 | 37243 | rs16866061 | 0.04363978 | 17834 | 78 |
| 2 | 233,559,301 | 233,753,501 | rs2675968 | 2.57E-06 | 621 | rs2675968 | 0.958718346 | 450 | 21 |
| 3 | 2,532,786 | 2,561,686 | rs17194476 | 7.29E-02 | 136349 | rs17620999 | 0.006866829 | 221634 | 33 |
| 3 | 17,221,366 | 17,888,266 | rs17044053 | 3.88E-03 | 14986 | rs11923589 | 0.063036738 | 10203 | 66 |
| 3 | 36,843,183 | 36,945,783 | rs4624519 | 6.26E-06 | 762 | rs4624519 | 0.838545989 | 698 | 12 |
| 3 | 52,541,105 | 52,903,405 | rs2239547 | 2.25E-06 | 610 | rs2239547 | 0.96441905 | 427 | 36 |
| 3 | 63,792,650 | 64,004,050 | rs832197 | 4.04E-04 | 3818 | rs11922435 | 0.257087548 | 2831 | 82 |
| 3 | 135,807,405 | 136,615,405 | rs10935182 | 1.23E-04 | 2154 | rs10935186 | 0.475527492 | 1548 | 39 |
| 3 | 180,588,843 | 181,205,585 | rs1351235 | 8.86E-06 | 836 | rs1805572 | 0.762139463 | 816 | 25 |
| 4 | 23,366,403 | 23,443,403 | rs215451 | 2.01E-03 | 9461 | rs215478 | 0.020263667 | 51863 | 92 |
| 4 | 103,146,888 | 103,198,090 | rs17823966 | 9.03E-03 | 27523 | rs170871 | 0.023889704 | 41691 | 6 |
| 4 | 170,357,552 | 170,646,052 | rs3797040 | 2.68E-04 | 3116 | rs3797040 | 0.433948011 | 1694 | 53 |
| 4 | 176,851,001 | 176,875,801 | rs1106568 | 4.31E-04 | 3933 | rs2333325 | 0.064505664 | 9890 | 76 |
| 5 | 45,291,475 | 45,393,775 | rs9292918 | 1.14E-02 | 32785 | rs6451798 | 0.009397056 | 143152 | 70 |
| 5 | 60,499,143 | 60,843,543 | rs34635 | 8.54E-05 | 1789 | rs7701440 | 0.544500484 | 1343 | 8 |
| 5 | 88,581,331 | 88,854,331 | rs187571 | 2.90E-02 | 66539 | rs187571 | 0.028878051 | 31739 | 65 |
| 5 | 109,030,036 | 109,209,066 | rs12656073 | 4.34E-03 | 16236 | rs12656073 | 0.085565957 | 7598 | 91 |
| 5 | 137,598,121 | 137,948,092 | rs13159624 | 2.38E-03 | 10684 | rs13159624 | 0.107687062 | 6135 | 67 |
| 5 | 140,023,664 | 140,222,664 | rs2337515 | 4.43E-03 | 16465 | rs17286731 | 0.080733657 | 8014 | 105 |
| 5 | 151,941,104 | 152,797,656 | rs12522297 | 7.23E-06 | 791 | rs2910032 | 0.093191565 | 7022 | 40 |
| 5 | 153,671,057 | 153,688,217 | rs6863455 | 4.31E-02 | 90406 | rs6863455 | 0.018396721 | 58777 | 93 |
| 6 | 26,000,000 | 34,000,000 | rs2021722 | 4.30E-11 | 1 | rs2021722 | 0.999990069 | 1 | 1 |
| 6 | 73,132,701 | 73,171,901 | rs9360557 | 1.01E-04 | 1936 | rs9360557 | 0.536308045 | 1367 | 89 |
| 6 | 84,279,922 | 84,407,274 | rs217297 | 7.86E-04 | 5388 | rs2224195 | 0.028036516 | 33156 | 47 |
| 6 | 96,300,000 | 96,500,000 | rs584453 | 2.35E-03 | 10575 | rs1546898 | 0.005915849 | 268799 | 54 |
| 7 | 1,896,096 | 2,190,096 | rs1107592 | 5.28E-08 | 141 | rs1107592 | 0.99673362 | 125 | 7 |
| 7 | 24,619,494 | 24,832,094 | rs2721783 | 5.60E-04 | 4505 | rs2711115 | 0.223822916 | 3219 | 90 |
| 7 | 86,403,226 | 86,459,326 | rs6943762 | 4.07E-03 | 15490 | rs12704289 | 0.024224333 | 40865 | 42 |
| 7 | 104,598,064 | 105,063,064 | rs4730073 | 9.81E-05 | 1900 | rs4730073 | 0.619997851 | 1105 | 50 |
| 7 | 110,034,393 | 110,106,693 | rs7783665 | 1.45E-04 | 2345 | rs7783185 | 0.080875631 | 7999 | 95 |
| 7 | 110,843,815 | 111,205,915 | rs38752 | 2.82E-04 | 3202 | rs214475 | 0.414654919 | 1784 | 15 |
| 7 | 131,539,263 | 131,567,263 | rs10954343 | 6.65E-04 | 4941 | rs10954343 | 0.069858026 | 9175 | 97 |
| 7 | 137,039,644 | 137,085,244 | rs320692 | 2.51E-04 | 3023 | rs320686 | 0.172770756 | 4030 | 60 |
| 8 | 4,177,794 | 4,192,544 | rs10503253 | 3.84E-07 | 333 | rs10503253 | 0.516813746 | 1404 | 77 |
| 8 | 27,412,627 | 27,453,627 | rs1565735 | 1.39E-03 | 7514 | rs7844965 | 0.043300639 | 18028 | 88 |
| 8 | 60,475,469 | 60,954,469 | rs7828435 | 9.82E-04 | 6114 | rs7838646 | 0.043872025 | 17687 | 71 |
| 8 | 89,340,626 | 89,753,626 | rs4484741 | 1.70E-07 | 216 | rs4484741 | 0.96417159 | 428 | 79 |
| 8 | 111,460,061 | 111,630,761 | rs17392088 | 3.91E-03 | 15059 | rs13270567 | 0.003609355 | 487064 | 32 |
| 8 | 143,309,503 | 143,330,533 | rs10098073 | 5.42E-06 | 736 | rs10098073 | 0.778332214 | 789 | 5 |
| 9 | 84,630,941 | 84,813,641 | rs2767713 | 7.53E-05 | 1694 | rs4877686 | 0.569316316 | 1270 | 61 |
| 10 | 18,681,005 | 18,770,105 | rs7893279 | 8.08E-04 | 5469 | rs7893279 | 0.028444957 | 32514 | 18 |
| 10 | 104,423,800 | 104,957,618 | rs11191580 | 2.23E-08 | 104 | rs11191580 | 0.998895413 | 69 | 3 |
| 11 | 24,367,320 | 24,412,990 | rs10834318 | 1.17E-04 | 2107 | rs10834318 | 0.231132207 | 3130 | 58 |
| 11 | 46,342,943 | 46,751,213 | rs12574668 | 8.95E-04 | 5801 | rs12574668 | 0.190473356 | 3717 | 24 |
| 11 | 57,386,294 | 57,682,294 | rs11570190 | 1.66E-05 | 974 | rs11570190 | 0.861473829 | 676 | 57 |
| 11 | 109,285,471 | 109,610,071 | rs1439513 | 1.69E-03 | 8492 | rs2212430 | 0.010916963 | 119181 | 94 |
| 11 | 113,317,794 | 113,423,994 | rs2514218 | 8.57E-03 | 26495 | rs4245147 | 0.025994085 | 36705 | 34 |
| 11 | 123,394,636 | 123,395,986 | rs7927176 | 8.42E-05 | 1772 | rs7927176 | 0.583233406 | 1228 | 73 |
| 11 | 124,610,007 | 124,620,147 | rs11219769 | 6.39E-02 | 122871 | rs11219769 | 0.022997524 | 44197 | 22 |
| 11 | 130,714,610 | 130,749,330 | rs10791097 | 2.17E-04 | 2853 | rs10791097 | 0.472430131 | 1555 | 16 |
| 11 | 133,808,069 | 133,852,969 | rs7106715 | 1.99E-04 | 2736 | rs7106715 | 0.267248486 | 2734 | 35 |
| 12 | 2,321,860 | 2,523,731 | rs4765905 | 8.92E-07 | 471 | rs4765905 | 0.980692241 | 321 | 4 |
| 12 | 29,905,265 | 29,940,365 | rs436124 | 1.91E-03 | 9158 | rs302321 | 0.136143977 | 5014 | 96 |
| 12 | 57,428,314 | 57,682,971 | rs324015 | 3.24E-04 | 3450 | rs324015 | 0.398537952 | 1857 | 19 |
| 12 | 92,243,186 | 92,258,286 | rs4240748 | 3.81E-03 | 14776 | rs4240748 | 0.019537175 | 54405 | 101 |
| 12 | 103,559,855 | 103,616,655 | rs998499 | 1.26E-02 | 35330 | rs10860949 | 0.004407257 | 394586 | 104 |
| 12 | 110,723,245 | 110,723,245 | rs4766428 | 1.93E-03 | 9249 | rs4766428 | 0.147415196 | 4640 | 52 |
| 12 | 123,448,113 | 123,909,113 | rs940904 | 4.93E-04 | 4211 | rs1727331 | 0.257456611 | 2825 | 9 |
| 14 | 30,189,985 | 30,190,316 | rs2068012 | 2.54E-02 | 60026 | rs2068012 | 0.03757315 | 22470 | 81 |
| 14 | 72,417,326 | 72,450,526 | rs12896825 | 2.80E-04 | 3191 | rs4902961 | 0.10553426 | 6258 | 69 |
| 14 | 99,707,919 | 99,719,219 | rs17098461 | 3.42E-01 | 490951 | rs17098461 | 0.008175246 | 176098 | 68 |
| 14 | 103,996,234 | 104,184,834 | rs4906356 | 8.41E-04 | 5586 | rs11846404 | 0.237856661 | 3040 | 13 |
| 15 | 40,566,759 | 40,602,237 | rs1869901 | 3.49E-07 | 323 | rs1869901 | 0.991093414 | 194 | 63 |
| 15 | 61,831,663 | 61,909,663 | rs4775413 | 4.01E-06 | 677 | rs11071612 | 0.297931316 | 2468 | 43 |
| 15 | 70,573,672 | 70,628,872 | rs1971791 | 3.27E-02 | 72789 | rs1971791 | 0.031307514 | 28593 | 86 |
| 15 | 78,803,032 | 78,859,610 | rs3813570 | 6.08E-04 | 4729 | rs3813570 | 0.285819187 | 2566 | 14 |
| 15 | 84,661,161 | 85,153,461 | rs11638630 | 4.59E-04 | 4061 | rs11631921 | 0.283185603 | 2588 | 28 |
| 15 | 91,416,560 | 91,429,040 | rs8032315 | 2.84E-03 | 12100 | rs8032315 | 0.114366082 | 5836 | 11 |
| 16 | 9,875,519 | 9,970,219 | rs9938117 | 6.22E-04 | 4776 | rs11647877 | 0.237327839 | 3046 | 80 |
| 16 | 13,728,459 | 13,761,359 | rs16962588 | 1.68E-03 | 8477 | rs16962588 | 0.000392095 | 892172 | 49 |
| 16 | 29,924,377 | 30,144,877 | rs12716974 | 2.20E-03 | 10117 | rs4283241 | 0.108968991 | 6075 | 37 |
| 16 | 58,669,293 | 58,682,833 | rs12447862 | 3.69E-02 | 80047 | rs12447862 | 0.010095183 | 131547 | 87 |
| 16 | 67,709,340 | 68,311,340 | rs2863981 | 3.28E-03 | 13303 | rs2863981 | 0.103787479 | 6354 | 83 |
| 17 | 2,095,899 | 2,220,799 | rs11078883 | 9.46E-04 | 5987 | rs11078883 | 0.228496217 | 3159 | 41 |
| 17 | 17,722,402 | 18,030,202 | rs2955384 | 5.23E-03 | 18490 | rs2955384 | 0.057142608 | 11854 | 85 |
| 18 | 52,747,686 | 53,200,117 | rs17512836 | 2.35E-08 | 106 | rs17512836 | 0.998854457 | 73 | 23 |
| 18 | 53,453,389 | 53,804,154 | rs17602354 | 9.33E-04 | 5943 | rs17602354 | 0.085071447 | 7632 | 29 |
| 19 | 19,374,022 | 19,658,022 | rs2965189 | 1.57E-04 | 2439 | rs2965189 | 0.533443545 | 1373 | 45 |
| 19 | 30,981,643 | 31,039,023 | rs919803 | 2.56E-02 | 60492 | rs919803 | 0.01109558 | 116888 | 64 |
| 19 | 50,067,499 | 50,135,399 | rs6509439 | 1.61E-04 | 2465 | rs11083979 | 0.355100419 | 2096 | 103 |
| 20 | 37,361,494 | 37,485,994 | rs4812319 | 3.69E-04 | 3671 | rs4812319 | 0.364286076 | 2046 | 26 |
| 20 | 48,114,136 | 48,131,649 | rs576119 | 1.85E-03 | 8994 | rs576119 | 0.135692494 | 5027 | 100 |
| 22 | 39,975,317 | 40,016,817 | rs5995756 | 3.67E-04 | 3661 | rs5995756 | 0.267076671 | 2735 | 38 |
| 22 | 41,408,556 | 41,675,156 | rs5758209 | 8.39E-06 | 826 | rs5758209 | 0.688851004 | 971 | 31 |
| 22 | 42,315,744 | 42,689,414 | rs134902 | 1.85E-04 | 2635 | rs134902 | 0.468962552 | 1568 | 55 |

<sup>a</sup> rank of the SNP with the smallest p-value
<sup>b</sup> rank of the SNP with the largest posterior score
<sup>c</sup> rank of 105 loci based on the p-values in the PGC2014 study

